# Whole organism and tissue specific analysis of pexophagy in *Drosophila*

**DOI:** 10.1101/2023.11.17.567516

**Authors:** Francesco G. Barone, Marco Marcello, Sylvie Urbé, Natalia Sanchez-Soriano, Michael J. Clague

## Abstract

Peroxisomes are essential organelles involved in critical metabolic processes in animals such as fatty acid oxidation, ether phospholipid production and reactive oxygen species detoxification. We have generated transgenic *Drosophila melanogaster* models expressing fluorescent reporters for the selective autophagy of peroxisomes, a process known as pexophagy. Using light sheet microscopy, we have been able to obtain a global overview of pexophagy levels, across the entire organism at different stages of development. Tissue specific control of pexophagy is exemplified by areas of peroxisome abundance but minimal pexophagy observed in clusters of oenocytes, which are the major site of long chain fatty acid synthesis. They are surrounded by epithelial cells where pexophagy is much more evident. Enhancement of pexophagy was achieved by feeding flies with the iron chelator deferiprone, in line with past results using mammalian cells. Specific drivers were used to visualise pexophagy in neurons, in which we tested the role of two proteins proposed to regulate pexophagy. Firstly, depletion of CG8814, the *Drosophila* homologue of the yeast protein Atg37, had no noticeable impact on pexophagy. In contrast, specific depletion in the larval central nervous system of Hsc70-5, the *Drosophila* homologue of the chaperone HSPA9/Mortalin, led to a substantial elevation in pexophagy.

## Introduction

Peroxisomes are discrete organelles that play key roles in lipid metabolism and reactive oxygen species (ROS) detoxification [1, 2]. They are also essential for cell death by the ferroptosis pathway [3, 4]. Their abundance is regulated via rates of synthesis and division on the one hand and selective degradation by autophagy on the other [5, 6]. This degradation process known as pexophagy can occur via different pathways, which involve either acquisition of BNIP3L (hereafter referred to as NIX) or the binding of ubiquitin receptors such as NBR1, both of which bind to LC3 on autophagosomal membranes [7-9]. Diseases linked to peroxisome homeostasis and function, known as peroxisome biogenesis or Zellweger spectrum disorders are profound multisystem diseases with severe effects on the central nervous system, for which it has been suggested that defective pexophagy is a fundamental feature [10, 11].

The distribution and size of peroxisomes within both cells and organisms has been studied by electron microscopy and more recently by expression of GFP-tagged peroxisomal markers [12, 13]. This has allowed depletion of peroxisomes to be linked to disease-associated mutations but does not formally distinguish between reduced biogenesis versus enhanced pexophagy. A fluorescence approach to measuring selective autophagy entails expressing a pH-sensitive ratiometric reporter protein that localises to the compartment in question. When these probes are delivered to acidic endolysosomal compartments, changes in their fluorescent properties can be visualised. For example two probes, mito-QC and mito-mKeima have been developed to measure the selective autophagy of mitochondria, known as mitophagy [14, 15]. We and others have substituted an SKL targeting signal to generate reporters for measuring pexophagy in mammalian cells [7, 8, 16, 17]. The ability to visualise mitophagy in intact organisms, such as mice or flies, has provided an insight into the tissue variation and changes during development. Analysis of mitophagy in PINK1 or Parkin mutant animals proved critical to realisation that the majority of basal mitophagy is independent of these linked genes [18, 19].

Peroxisomes in *Drosophila* have been analysed using expression of GFP-reporters in genetic screens of peroxisomal size or number [12, 20, 21]. Here we have extended this approach to directly visualise the magnitude of pexophagy alongside these parameters, by adopting both pexo-QC and pexo-Keima reporters. Furthermore, we take advantage of light sheet microscopy to capture the distribution of pexophagy across developmental stages of the whole organism in a single image. As a first application of these models we have performed imaging experiments on whole larvae (*in vivo*) and adult brains (*ex vivo*) to examine the impact of two proteins with proposed roles in regulating pexophagy; CG8814, the *Drosophila* homologue of Atg37/ACBD5 and secondly the Hsc70-5 (Mortalin) gene [21, 22].

## Results

### Characterisation of pexophagy reporter flies

We started by making Gal4/UAS-inducible transgenic *Drosophila* strains that express either EGFP-mCherry-SKL (hereafter pexo-QC) or Keima-SKL (hereafter pexo-Keima) pexophagy reporters, which were chosen for their ability to detect pH changes within autophagic structures (Figure 1 A, B and S1A). For each reporter, multiple lines with equivalent expression were developed and subsequently a single line was selected for further analysis. We observed that the ubiquitous expression of these reporters, using a Tubulin Gal4 driver (tub-Gal4), had no apparent impact on development or viability, suggesting that they were well tolerated by the organism. To verify the expected targeting to peroxisomes we examined colocalisation with the peroxisomal membrane protein PMP34 in larval epidermis. We found that both pexo-QC and pexo-Keima reporter colocalise well with peroxisomes (PMP34-Cer positive organelles) in the green pH-neutral channel as expected (Figure 1 C and Figure S1B). First indications of pexophagy are evidenced by magenta punctae (derived from pexo-QC) that colocalise with the marker of acidic endolysosomal compartments, LysoTracker™ (Figure 1D). Similarly, pexo-Keima’s fluorescence excitation spectrum shifts at low pH, with an enhanced signal from excitation at 561 nm (Figure S1A). We next quantitated the number of pexolysomes per cell following two experimental interventions. The iron chelator deferiprone (DFP) is known to induce pexophagy in mammalian cells using the pexo-QC reporter [8, 9]. Here we show that its administration to flies, via a feeding regimen, produces a similar effect (Figure 1E,F). We also expressed an RNAi transgene suppressing the core component of the autophagy machinery, Atg5, which strongly reduced the pexophagy levels (Figure 1E,F). Qualitatively comparable results were found with pexo-Keima flies (Figure S1C). In all subsequent experiments we have used pexo-QC expression as it showed superior brightness and has the advantage of being fixable.

**Figure 1:**
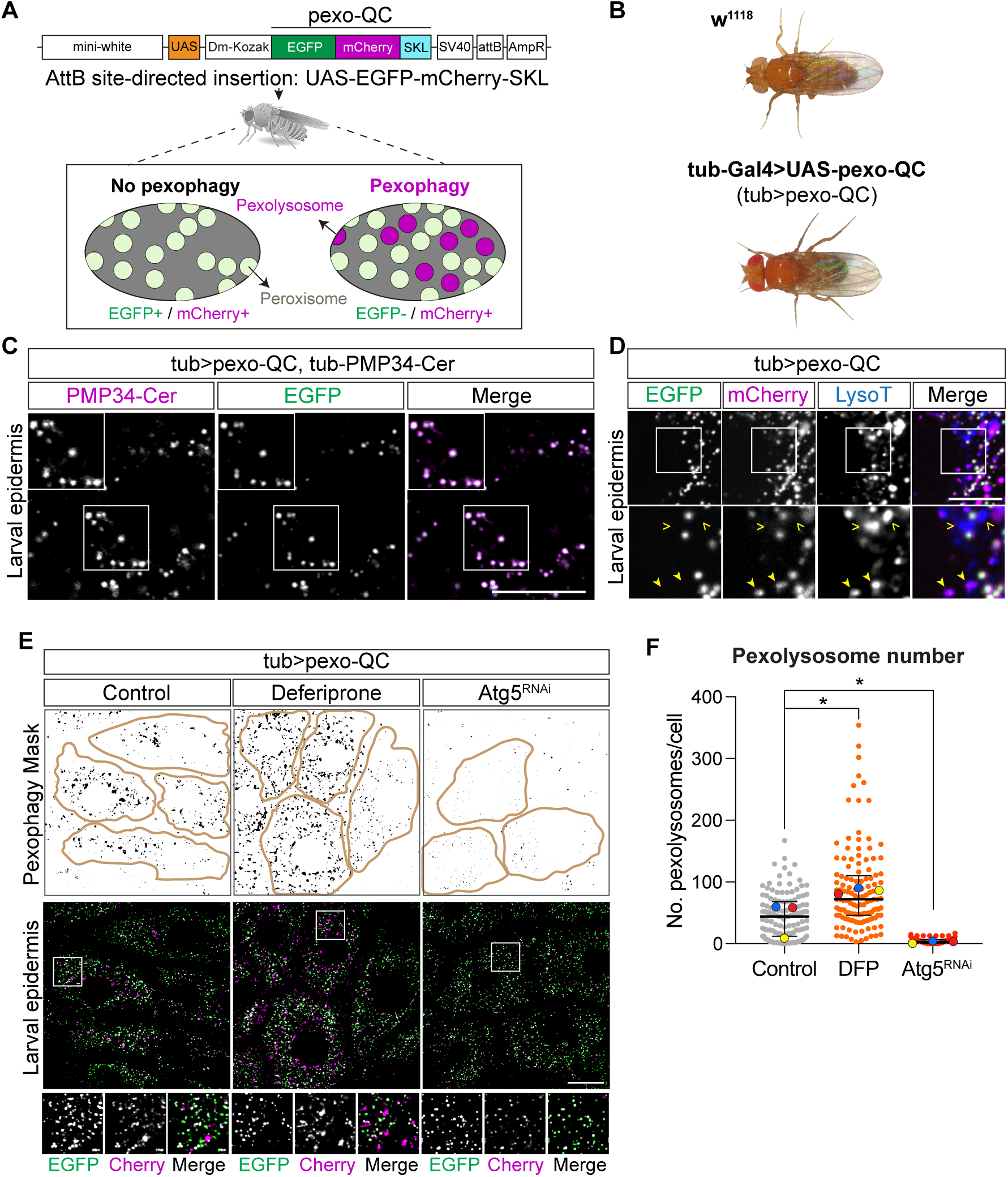
Generation of the pexo-QC *Drosophila melanogaster* for *in vivo* detection of pexophagy and peroxisomal abundance. (**A**) Diagram of the cassette used for expression of a EGFP-mCherry-SKL fusion protein (pexo-QC) used to create the pexo-QC *Drosophila* (elements not to scale). This includes a mini-white gene, upstream activating sequence (UAS), a *Drosophila* melanogaster Kozak sequence (Dm-Kozak), the EGFP-mCherry-SKL coding sequence, an SV40 poly adenylation sequence, an attB site for site-directed insertion, and an ampicillin resistance gene (AmpR). At neutral pH, peroxisomes display both GFP and Cherry fluorescence whereas only Cherry fluorescence is retained at the acidic pH of the lysosome (pexolysosomes, magenta) (**B**) Bright field images of w^1118^ flies and tub-Gal4>UAS-pexo-QC flies (9-12 days old), indicating no observable phenotypical changes or gross morphological abnormalities in the pexo-QC *Drosophila* model. (**C**) High-resolution Airyscan live images of L3 larval epidermis comparing EGFP fluorescence signal (green) of pexo-QC with exogenous expression of peroxisomal membrane protein PMP34-Cer (Tub-PMP34-Cerulean, shown in magenta). Representative of three independent experiments, with at least three animals per condition in each experiment. Scale bar: 20 μm. (**D**) Confocal live imaging analysis of pexo-QC larval epidermal cells costained with LysoTracker^TM^ to identify lysosomes (blue). Representative of three independent experiments, with at least three animals per condition in each experiment. > LysoTracker^(+)^, mCherry^(-)^ puncta; filled arrowhead, LysoTracker^(+)^/ mCherry^(+)^ puncta. Scale bar: 10 μm (**E**) Representative confocal images of pexo-QC larval epidermal cells upon expression of an Atg5-RNAi transgene (Atg5^RNAi^) to inhibit autophagy or exposure to the iron chelator deferiprone (DFP, 65 *µ*M) to induce pexophagy. Scale bar: 20 μm. Also shown is a pexophagy mask, automatically generated from the mCherry/GFP ratio using the Fiji macro previously reported [53]. (**F**) Graph shows the number of pexolysosomes per cell for each condition. The mean and SD of three colour-coded independent experiments are shown; one-way ANOVA with Dunnett’s multiple comparison test for three colour-coded independent experiments with a minimum of three animals per condition in each experiment. *P < 0.05. Genotypes analysed were *w^1118^; UAS-pexo-QC/+; tub-Gal4/tub-PMP34-Cer (C), w^1118^; UAS-pexo-QC/+; tub-Gal4/+ (D), and w^1118^; UAS-pexo-QC/+; tub-Gal4/+, w^1118^; UAS-pexo-QC/UAS-Atg5^RNAi^; tub-Gal4/+ (E)*.

### A whole organism view of pexophagy gained via light sheet microscopy

We employed light sheet microscopy, alongside spinning disk confocal microscopy, to visualise the whole organism at different stages of development, using the Tubulin Gal4 driver for ubiquitous pexo-QC expression. Pexophagy was evident across the organism, but unevenly distributed, with clear hot spots discernible by confocal microscopy in the living late stage embryo (Figure S2). We also employed fixation prior to visualisation of third instar larvae (hereafter L3) and white pre-pupae by light sheet microscopy (Figure 2 and S3, S4). Highly elevated areas of pexophagy are clearly evident within the central nervous system, midgut and anal plate (Figure 2B-F). Overall, pexophagy is most prominent in the pre-pupae (Figure 2G). There are also clear areas rich in peroxisomes but low in pexophagy including seven peroxisome rich, pexophagy poor clusters comprising 4-6 cells located in a basal position below the epidermis (Figure 3A-C). These represent oenocytes, which are large secretory, hepatocyte-like cells in *Drosophila* and the major sites for very long chain fatty acid synthesis [23]. In line with this function, they are known to be sites of peroxisome enrichment [24]. Our data clearly indicate that peroxisomes exhibit a very low turnover rate in this organ, which we propose underlies their enrichment. In contrast, oenocytes are surrounded by epidermal cells, which show significantly higher levels of pexophagy (Figure 3B).

**Figure 2:**
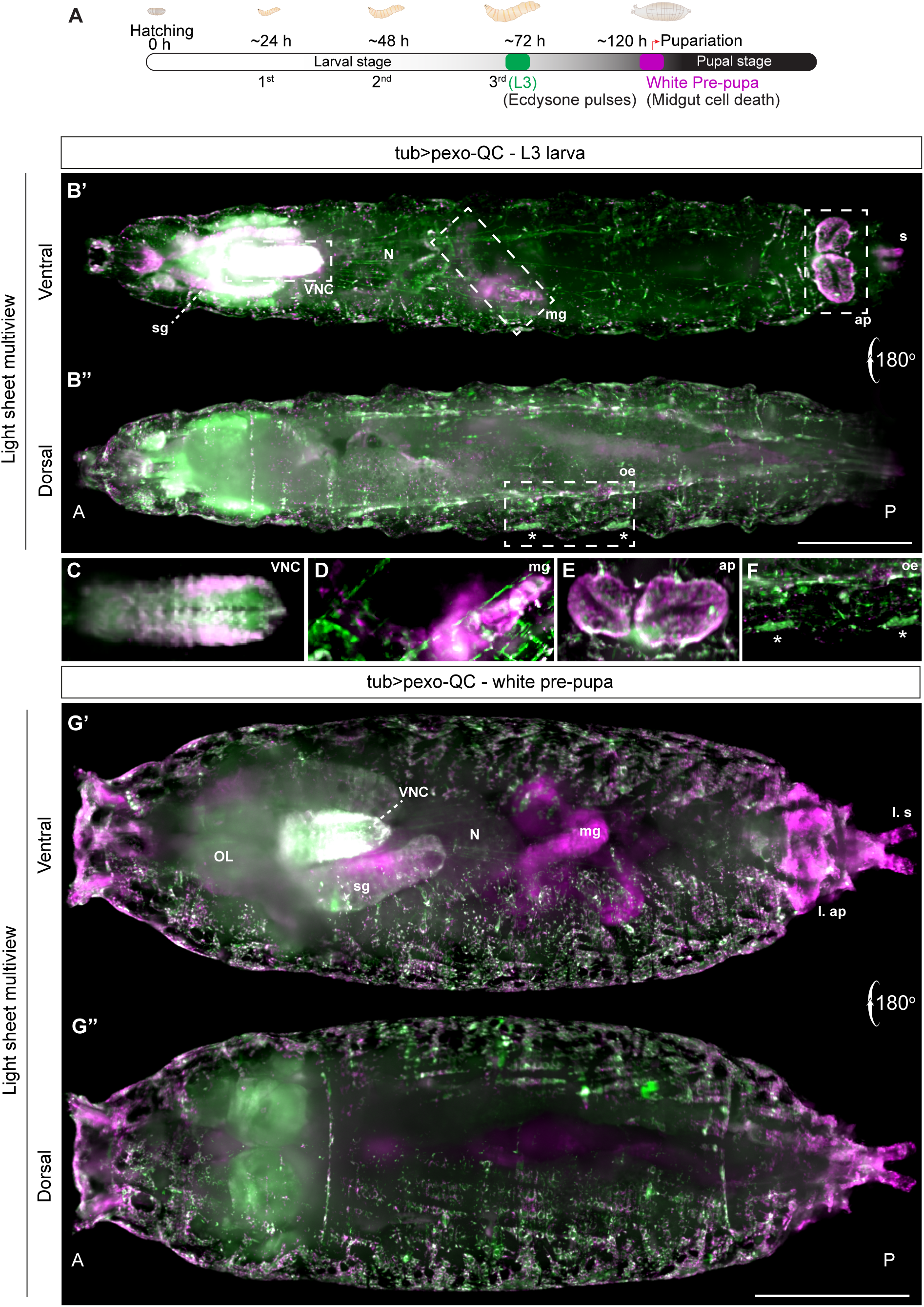
Light sheet microscopy reveals tissue-specific distribution of *in vivo* pexophagy in intact *Drosophila* larvae and prepupa. (**A**) Schematic representation of *Drosophila* embryo, larval development and early pupariation stage (white pre-pupa). (**B’-B’’**) Light Sheet Fluorescence Microscopy (LSFM) multiview images of pexo-QC L3 larva (tub>pexo-QC), ventral view B’, dorsal view B’’ (**C-D-E-F**) Enlarged insets of the specimen shown in B’ and B’’. Z-thickness and image contrast settings were adjusted to highlight the differential distribution of peroxisomes and pexolysosomes in the respective tissues. (**C**) Pexophagy in the Ventral Nerve Chord (VNC) with heterogeneous pexophagy levels. (**D**) Enrichment of pexolysosomes in a portion of the larval midgut (mg). (**E**) Enrichment of pexolysosomes in the larval anal plates (ap). Insets shown in D and E were rotated by -45° and -90° respectively for improved visualisation and clarity. (**F**) Peroxisomal enrichment in larval oenocyte (oe) clusters. Asterisks indicate two individual oenocyte clusters. Salivary glands (sg); Nerves (N). (**G’-G’’**) LSFM multiview images of white pexo-QC pre-pupae (tub>pexo-QC, ventral view G’, dorsal view G’’). Optical lobes (OL); Salivary glands (sg); Larval anal plates (L. ap); Larval spiracles (L. s). The genotype analysed was *w^1118^; UAS-pexo-QC/+; tub-Gal4/+*. Scale Bar 500 μm. Single channel images of B’, B” and G’, G” are shown in Figure S3 and S4 respectively.

**Figure 3:**
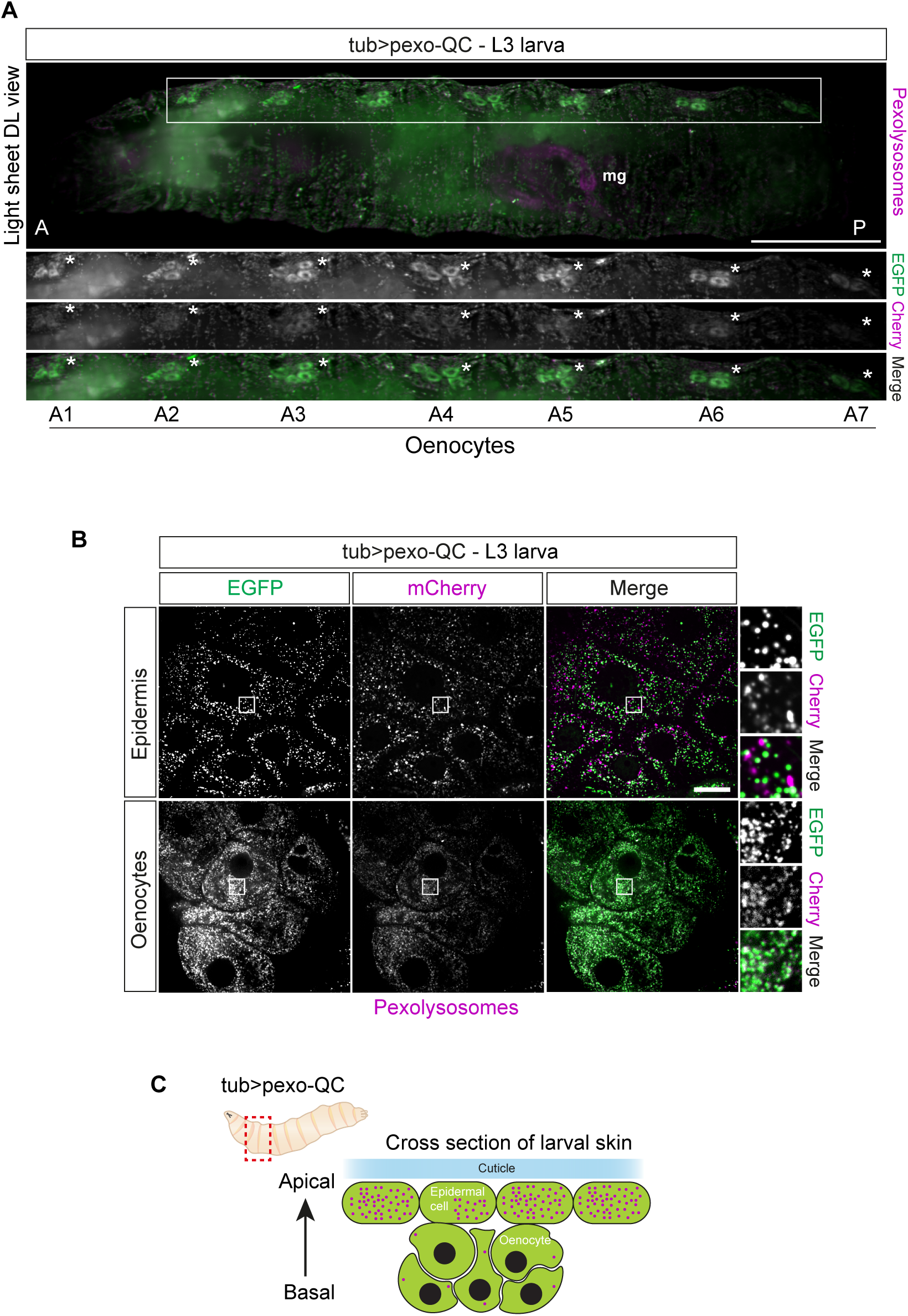
*Drosophila* oenocytes are enriched with peroxisomes but show low levels of pexophagy. (**A**) A representative Z-stack maximum intensity projection image acquired from a dorsolateral (DL) view of pexo-QC L3 larva (tub>pexo-QC) using a Light sheet microscope reveals seven identifiable peroxisome rich clusters (white asterisks, A1-A7) that based on their localisation and morphology can be classified as oenocytes. Scale bar: 500 μm. (**B**) Representative confocal images comparing pexophagy levels between oenocytes and the surrounding epidermis in pexo-QC larvae using the same driver (tub-Gal4). Images were acquired using a 3i confocal microscope with a 63x NA 1.4 objective lens. Scale bar: 20 μm. (**C**) Schematic representation of a cross section of the larval skin in *Drosophila melanogaster*, showing the relative positions of epidermal cells, rich in pexolysosomes, (shown in magenta) and oenocytes which have few pexolysosomes. Genotype analysed was *w^1118^; UAS-pexo-QC / +; tub-Gal4/+*.

### Visualisation of Pexophagy within the central and peripheral nervous system

The sheer density and complexity of the central nervous system requires specific drivers to visualise different cell populations *in vivo*. We have separately expressed pexo-QC under the control of pan-neuronal (elav-Gal4), motor neuronal (ok6-Gal4) and pan-glial (repo-Gal4) drivers (Figure 4A,B). All three specific drivers show enhanced pexophagy in clusters close to the ventral midline. It is clear that at the 3rd instar larval stage, the cell bodies of motor neurons show much higher levels of pexophagy than the total neuronal population, with high pexophagy apparent along the midline (Figure 4C,D). In addition, in the peripheral nervous system, individual peroxisomes can be visualised in the axonal projections of motor neurons, which exhibit a high degree of pexophagy. In the glia, surrounding the nerves, the density of peroxisomes is found to be higher and the pexophagy index correspondingly lower (Figure 4E). For analysis of the adult *Drosophila* brain it was necessary to perform *ex vivo* experiments. We found pexophagy to be prominent in the optic lobe (neurons and glia) and for motor neurons, in the suboesophageal zone (SEZ) (Figure 5).

**Figure 4:**
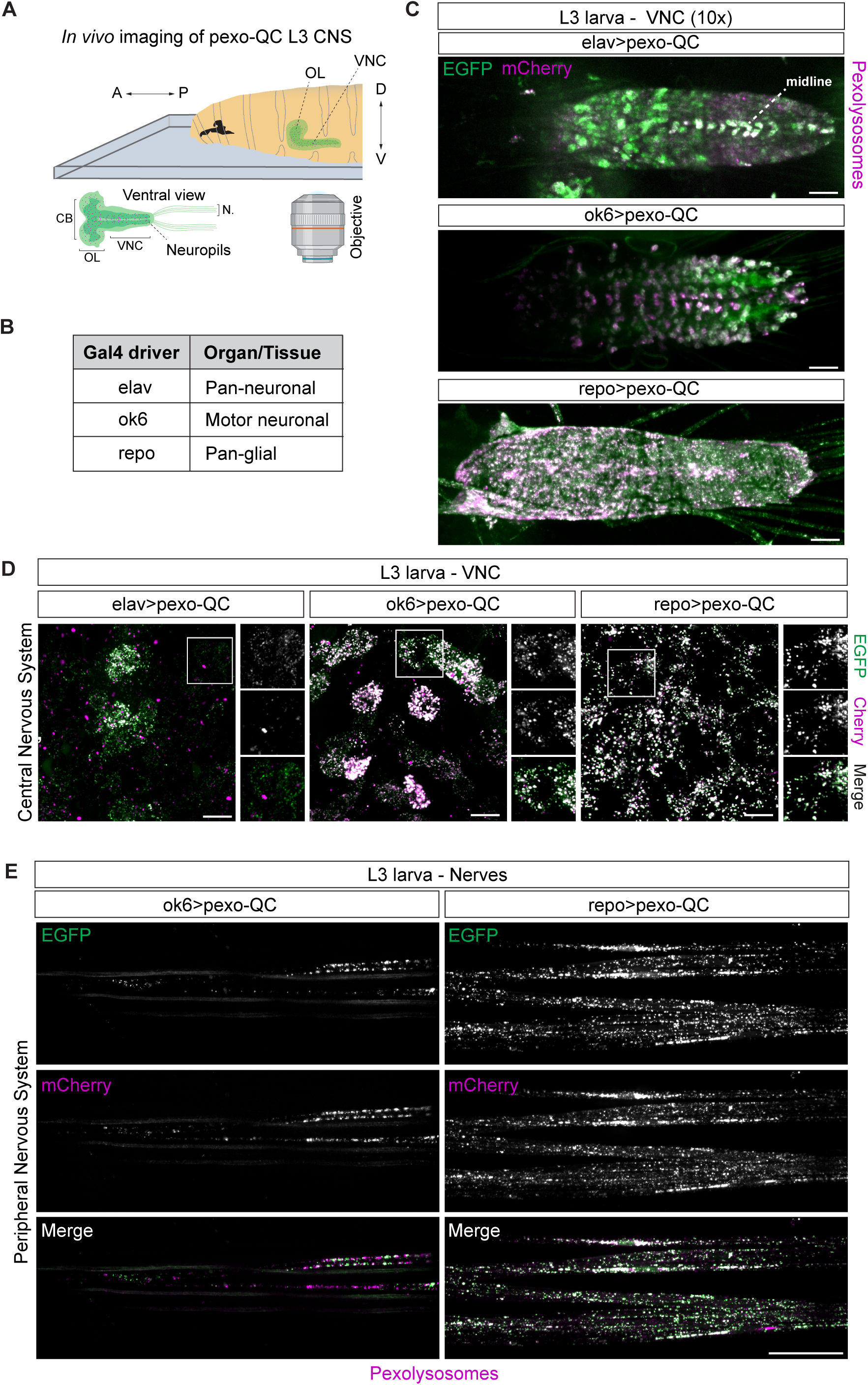
Visualisation of neuronal and glia pexophagy (*in vivo*) in the nervous system of *Drosophila* Larvae. (**A**) The imaging setup utilised to explore *in vivo* pexophagy in the nervous system of pexo-QC L3 larva using a 3i spinning disc confocal microscope is depicted schematically. Top: Position of larvae immobilised in 10% chloroform halocarbon oil. Bottom: ventral view of the sample’s central and peripheral neural systems, Ventral Nerve Cord (VNC) and nerves (N), respectively. A: Anterior; P: Posterior; D: Dorsal; V: Ventral; CB: Central Brain; OL: optical lobes. (**B**) Table displaying the different neural drivers employed in the study to analyse pexophagy in the pexo-QC larvae’s nervous system: elav-Gal4, a pan-neuronal, ok6-Gal4, a motor neuronal and repo-Gal4, a pan-glial driver. (**C**) Representative maximum intensity projection images of confocal Z-stacks of the VNC of pexo-QC larvae acquired using the imaging setup described in (A). Images were acquired using a 10x NA 0.3 objective. Scale bar: 50 μm. (**D**) Higher magnification images of maximum projection images of confocal Z-stacks of the VNC using a 63x NA 1.4 oil immersion objective. Scale bar: 10 μm. (**E**) Representative confocal maximum intensity projection images of confocal Z-stacks of nerves using distinct neuronal drivers. Left. *In vivo* pexophagy in motor neurons (ok6-Gal4). Right. *In vivo* pexophagy in wrapping glial cells (repo-Gal4). Scale bar: 20 μm. Genotypes analysed were *w^1118^; UAS-pexo-QC/+; elav-Gal4/+* (neurons), *w^1118^; UAS-pexo-QC/ok6-Gal4; +/+* (motor neurons), *w^1118^; UAS-pexo-QC/+; repo-Gal4/+* (glial cells).

**Figure 5:**
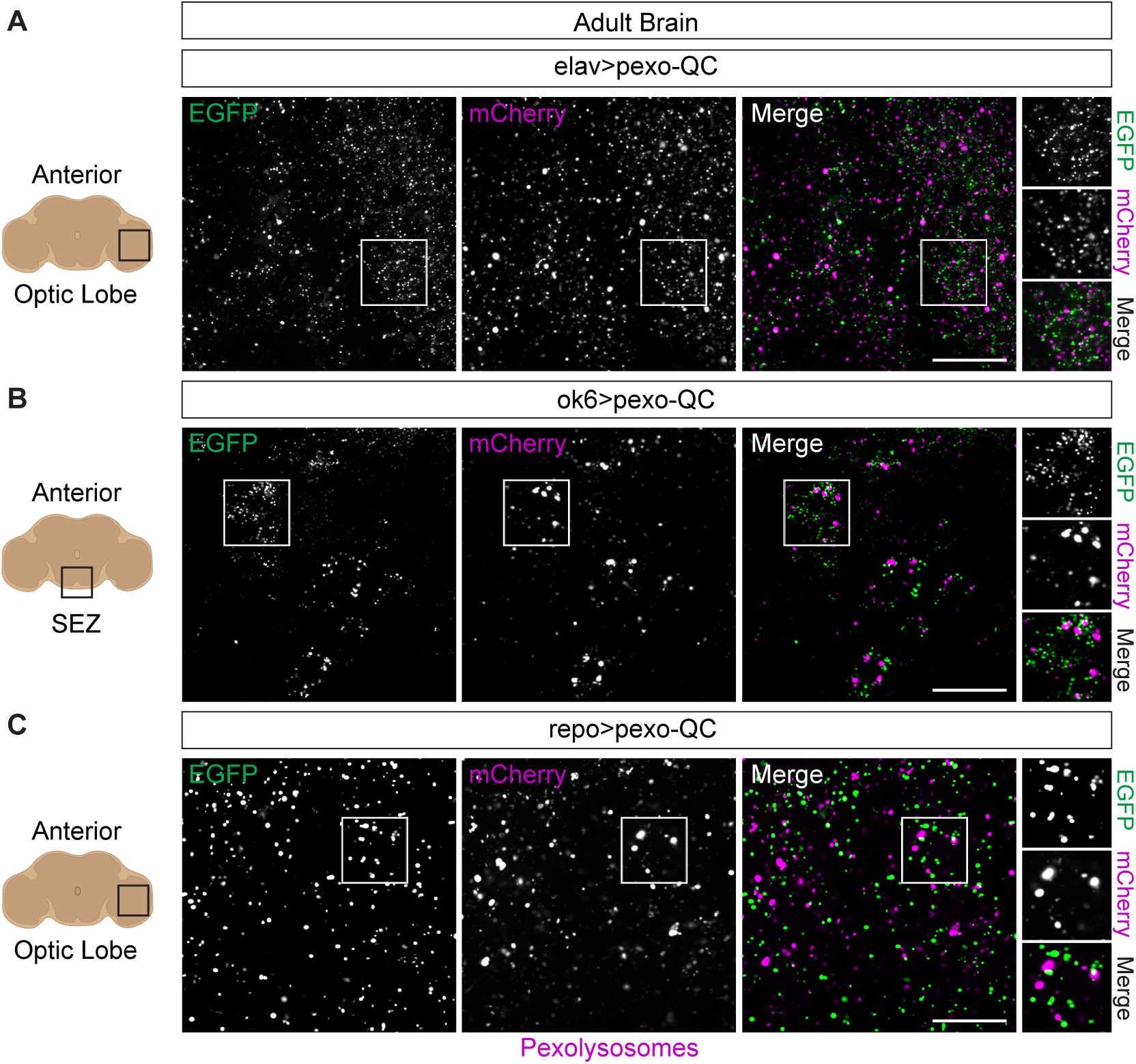
Pexophagy (*ex vivo) in* neuronal and glial cells in the adult central nervous system of pexo-QC flies. Representative confocal images of pexo-QC adult brain from 9-12 days old flies, using distinct neural drivers (3i spinning disc confocal microscope, 63x NA 1.4 objective). (**A**) Neuronal pexophagy in the optical lobe of *ex vivo Drosophila* pexo-QC adult brain (elav>pexo-QC). (**B**) *Ex vivo* pexophagy in the motor neurons of the suboesophageal zone (SEZ) of *Drosophila* pexo-QC adult brain (ok6>pexo-QC). (**C**) Glial pexophagy in the optical lobe of *ex vivo Drosophila* pexo-QC adult brain (repo>pexo-QC). Genotypes analysed were *w^1118^; UAS-pexo-QC/+; elav-Gal4/+* (neurons, A), *w^1118^; UAS-pexo-QC/ok6-Gal4; +/+* (motor neurons, B), *w^1118^; UAS-pexo-QC/+; repo-Gal4/+* (glial cells, C). Scale bar: 20 μm.

### Direct testing of gene function in the pexophagy pathway

ACBD4 and ACBD5 are members of the acyl-CoA binding domain (ACBD) protein family in humans that are primarily involved in lipid metabolism and vesicular trafficking [25]. Mutations in the *ACBD5* gene have also been associated with several human conditions, including peroxisome disorders and neurological disease [26-28]. Both of these proteins contain a highly conserved transmembrane domain required for peroxisomal localisation [29]. Depletion of the yeast ACBD5 homologue, ATG37, has been shown to reduce pexophagy. Similar claims have been made for ACBD5 in mammalian cells but this has not been supported by our recent data [9, 30]. CG8814 is the unique *Drosophila* orthologue of ACBD4 and ACBD5, hereafter referred to as Acbd4-5 (Figure S5A). A tagged version, Acbd4-5-3xHA localises to the periphery of peroxisomes in L3 larvae as evidenced by co-localisation with pexo-QC (Figure S5B). We next investigated the impact on pexophagy of reducing Acbd4-5 by RNAi. An Acbd4-5 targeting RNAi line was acquired and crossed with the pexo-QC model flies expressed either by (ubiquitous) tub or pan-neuronal elav drivers. In parallel, we also used two Acbd4-5 KO alleles (Acbd4-5^SK3^ and Acbd4-5^SK6^) in heterozygosis, in combination with pexo-QC in neurons (elav-driver). In no case could we detect any change in pexophagy parameters in either larval epidermis (RNAi, tub driver), larval nervous system (RNAi, elav driver) or adult central nervous system (RNAi, Acbd4-5^SK3^ and Acbd4-5^SK6^ and elav driver), whilst mRNA levels are reduced by more than 50%. (Figure 6A-H and S6).

**Figure 6:**
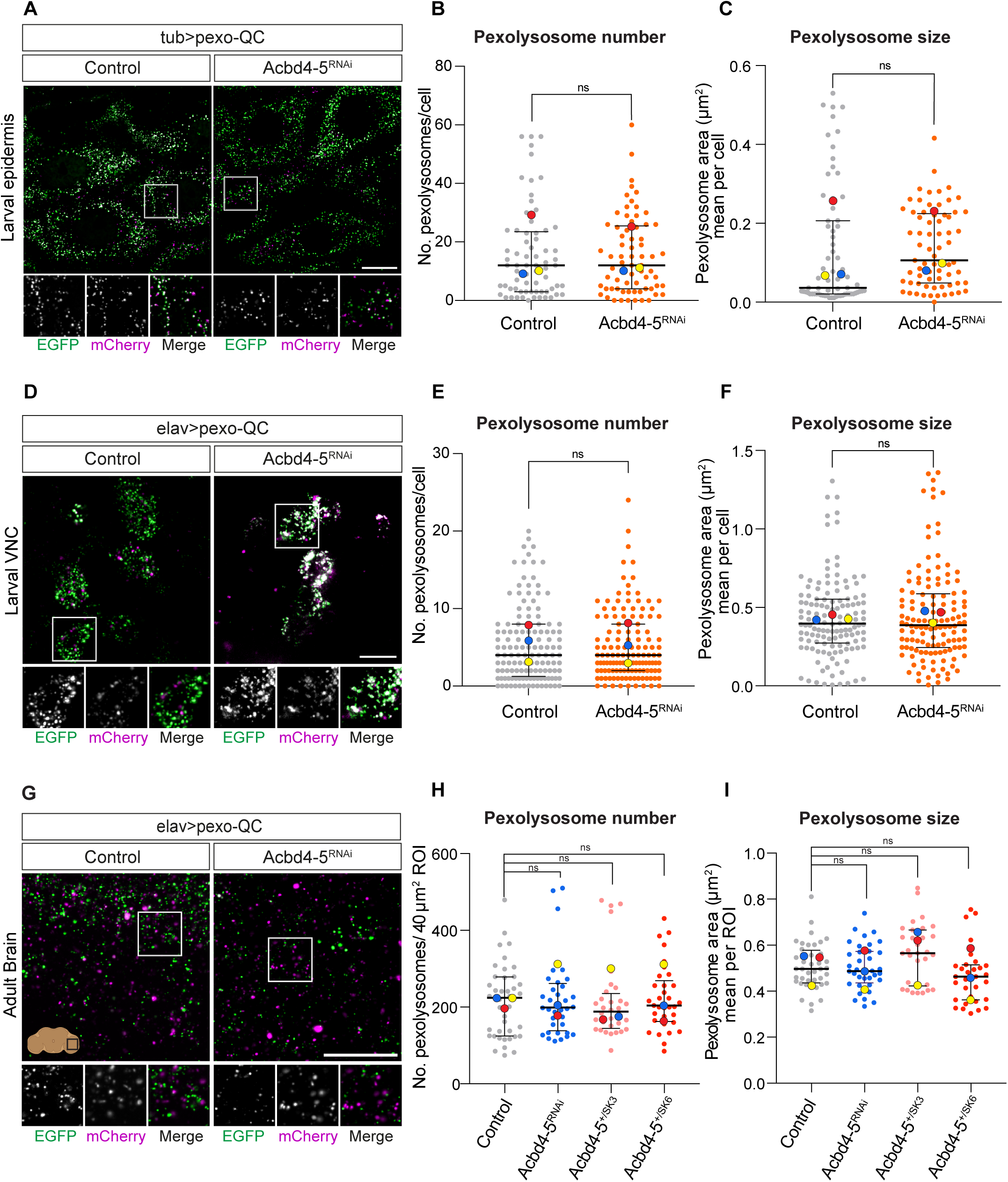
Depletion of Acbd4-5 does not affect neuronal pexophagy in pexo-QC larvae and adult brains. (**A**) Representative confocal images of *in vivo* larval epidermal cells (tub>pexo-QC) ± Acbd4-5 depletion (Acbd4-5^RNAi^). Scale bar: 20 μm (**B-C**) Graphs shows the number and mean area (size) of pexolysosomes per cell for each condition. At least 3 animals per condition in each experiment were analysed and >20 cells per condition quantified in three independent replicate experiments. Statistical significance determined by unpaired t-test; ns = not significant. The mean and SD of three colour-coded independent experiments are shown. (**D**) Representative Z-stack maximum intensity projection images of *in vivo* larval VNC neurons (elav>pexo-QC) ± Acbd4-5 depletion (Acbd4-5^RNAi^). Scale bar: 10 μm.(**E-F**) Graphs indicate the total number of pexolysosomes and their mean size in each cell. At least 3 animals per condition in each experiment were analysed and >40 cells were quantified per condition in three replicate experiments. Statistical significance determined by unpaired t-test; ns = not significant. The mean and SD of three colour-coded independent experiments are shown. (**G**) Representative maximum intensity projection of a Z-stack of confocal images of *ex vivo* adult brain (9-12 days old) neurons (elav>pexo-QC) ± Acbd4-5 depletion (Acbd4-5^RNAi^). (**H-I**) Graphs illustrate the number and mean area of pexolysosomes per 40 μm^2^ region of interest (ROI) of controls, Acbd4-5^RNAi^ and Acbd4-5^SK3^ and Acbd4-5^SK6^ KO alleles in heterozygosis (see Figure S6 for the representative images of the Acbd4-5^SK3^ and Acbd4-5^SK6^ heterozygotes). At least 9 ROI were quantified per condition in each replicate experiment from at least 3 animals. The mean and SD of three colour-coded independent experiments are shown. One-way ANOVA with Dunnett’s multiple comparison test; ns = not significant. Scale bar: 20 μm. Genotypes analysed were *w^1118^; UAS-pexo-QC/+; tub-Gal4/+, w^1118^; UAS-pexo-QC/UAS-Acbd4-5^RNAi^; tub-Gal4/+, w^1118^; UAS-pexo-QC/+; elav-Gal4/+, w^1118^; UAS-pexo-QC/UAS-Acbd4-5^RNAi^; elav-Gal4/+, w^1118^; UAS-pexo-QC/Acbd4-5^SK3^; elav-Gal4/+, w^1118^; UAS-pexo-QC/Acbd4-5^SK6^; elav-Gal4/+*.

Mortalin/HSPA9 was identified as a pexophagy regulator and shown to partially localise to peroxisomes [21]. Reduction of this protein has been linked to neurodegenerative diseases such as Parkinson’s and Alzheimer’s [31-33]. Jo et al. have shown a loss of GFP-tagged peroxisomes in flies following depletion of the Mortalin *Drosophila* homologue Hsc70-5, but did not directly visualise pexophagy, nor investigate its role in the central nervous system [21]. Here we use *in vivo* imaging of pexo-QC to confirm a strong increase in pexophagy in the larval ventral nerve cord following Hsc70-5 depletion with a previously validated RNAi line against Hsc70-5 (UAS-Hsc70-5^RNAi^, [21]) (Figure 7). This data confirms that *Drosophila* Hsc70-5 is a suppressor of pexophagy and validates our pexo-QC line as an efficient tool to measure pexophagy.

**Figure 7:**
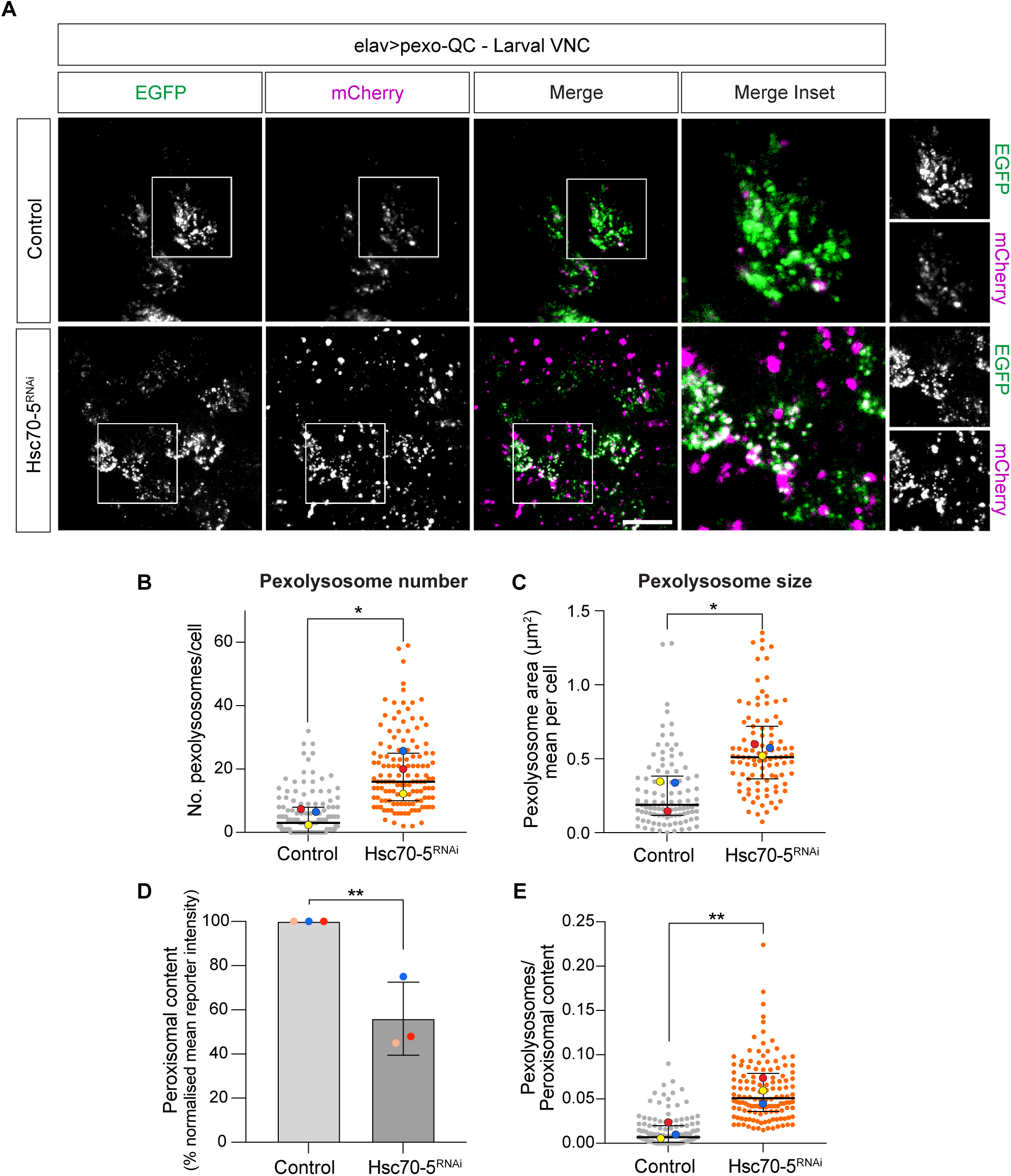
Hcs70-5 depletion enhances *in vivo* basal pexophagy in the ventral nerve chord of L3 larvae. (**A**) Representative Z-stack maximum intensity projection images of *in vivo* larval VNC neurons (elav>pexo-QC) ± Hcs70-5 depletion (Hcs70-5^RNAi^). (**B-E**) Graphs show the number and mean area (size) of pexolysosomes per cell as well as peroxisomal content (percentage of normalised mean reporter intensity, EGFP) and the pexolysosome to peroxisomal content ratio. At least 30 cells from a minimum of three animals were analysed per condition in each independent experiment. The mean and SD of three colour-coded independent experiments are indicated. Statistical significance was determined by unpaired t-test; \**P* < 0.05, \*\**P* < 0.01. Scale bars 10 μm. Genotypes analysed were *w^1118^; UAS-pexo-QC/+; elav-Gal4/+, w^1118^; UAS-pexo-QC/UAS-Hcs70-5^RNAi^; elav-Gal4/+*.

## Discussion

The introduction of selective autophagy reporters into whole organismal models has been applied to nematodes, flies, zebrafish and mice [15, 18, 34-36]. Each organism offers different pros and cons. For flies, the advantages include a relatively simple anatomy and low husbandry costs. Here, we also highlight their suitability for light sheet imaging of the entire fly larvae and pre-pupae, providing the first whole organism overview of any form of selective autophagy.

Mitophagy has received particular attention and its visualisation combined with genetic manipulation has allowed assessment of the contribution of different molecular mechanisms in the context of different organisms [18, 19, 34]. For example, the PINK1/Parkin pathway has been definitively shown to contribute little to the bulk turnover of mitochondria in flies and mice [18, 19]. Peroxisomes and mitochondria cooperate in the metabolism of cellular lipids and reactive oxygen species, in fact sharing many proteins [37]. Both pexophagy and mitophagy may proceed through a NIX/BNIP3-dependent pathway which links the respective membranes to autophagosomes [8, 9, 38-40]. It has been proposed that their turnover may be coupled via regulation of NIX, but different modes of NIX regulation at each organelle can exist. We envision a future study wherein our pexo-QC fly can be directly compared with the mito-QC model to investigate this issue.

The introduction of any new fly reporter model enables a genetic strategy to test literature claims of gene function. Here, our data do not support a significant role of the *Drosophila* ACBD5 homologue in pexophagy. This lies contrary to studies in yeast (Atg37), but chimes with our recent studies in mammalian cells [9, 22, 41]. We turned to a second opportunity to showcase the ability of our model to reflect changes in pexophagy. Our finding that reduction in Hsc70-5 enhances pexophagy confirms the model of Jo et al., who used indirect assays to infer this effect [21]. Thus, in the present study, we have demonstrated both chemical (DFP) and genetic enhancement of pexophagy.

The ability to identify tissues showing distinct pexophagy profiles, within an organism, is exemplified by our finding that oenocytes show extremely low levels of pexophagy, whilst being replete with peroxisomes. As peroxisomes play especially important roles in this tissue, to effect very long chain fatty acid synthesis, it is interesting to note their correspondingly privileged protection from turnover [23]. One possible mechanism would be suppression of the fly homologue of NIX/BNIP3, CG5059, but this is not imposed at the transcriptional level, according to the Fly Cell Atlas project data [8, 9, 42].

In summary, we introduce a tool for pexophagy research that will provide the field with new opportunities to assess gene function and metabolic control mechanisms. Our flies add to the suite of organismal models for selective autophagy and open up opportunities for direct comparison with those that use the same reporter type (organelle-QC).

## Acknowledgements

FB has been funded by a Wellcome Trust PhD studentship (102172/B/13/Z). MC is a Royal Society Industry Fellow (INF\R2\212031). NSS is supported by the BBSRC (BB/R018960/1 and BB/W016907/1). We thank the Liverpool Centre for Cell Imaging for providing access to facilities and Emma Rusilowicz-Jones for help with maintenance of fly stocks, the *Drosophila* fly facility, University of Manchester, especially Sanjai Patel for outstanding support and generation of transgenic flies. Stocks obtained from the Bloomington *Drosophila* Stock Center (NIH P40OD018537), the Vienna Drosophila Resource Center (VDRC), FlyORF (Zurich) and the Fly Stock National Institute of Genetics (NIG-FLY, Japan) were used in this study.

## Material and Methods

### Drosophila stocks, husbandry, and reagents

Flies were maintained on standard fly food at 25°C in the common cornmeal-agar medium. Fly stocks were obtained from BDSC (indicated with BL#), VDRC, FlyORF (indicated with F#), NIG, or as otherwise indicated. Stock used in this study are: w^1118^ [43] and tub-Gal4 III [44], ok6-Gal4 II (a gift from C. O’Kane [45]), Repo-Gal4 III [46], elav-Gal4 III [47], UAS-atg5 RNAi II (VDRC, 104461), UAS-pexo-QC II (this study), UAS-Keima-SKL II (this study), UAS-Hsc70-5 RNAi KK (VDRC, KK106236), UAS-Acbd4-5 RNAi II (CG8814, BL 67020), UAS-Acbd4-5-ORF.3xHA III (CG8814, F002670), Acbd4-5^SK3^ (CG8814, NIG, M2L-1612), Acbd4-5^SK6^ (CG8814, NIG, M2L-1613), tub-PMP34-Cerulean III (BL64246). All larvae shown in this study were actively feeding at time of collection, unless otherwise indicated. To administer DFP, pexo-QC and pexo-Keima (mKeima-SKL) L2 Larvae were starved for 4 hours and then transferred to a tube containing a mixture of water, 5% sucrose, and 65 *µ*M DFP. After a 20-hour incubation period, 3^rd^ instar larvae (L3) were selected and imaged to measure pexophagy levels. Drug administration in *Drosophila* larvae and DFP concentration adapted from Kim et al. and Lee et al respectively [18, 48]. All reagents were purchased from Sigma Aldrich unless otherwise stated.

### Transgenic line construction

A pexo-QC construct (pcDNA3.1-EGFP-mCherry-SKL was generated by PCR amplifying pEGFP from pEGFP-C1 and inserting this into pcDNA3.1-mCherry-SKL (addgene, #54520). pcDNA3.1-EGFP-mCherry-SKL was inserted in a pUAST.attB (DGRC, #1419) destination vector, for specific phiC31-based transgenesis. For the pexo-Keima line, Keima-SKL was PCR-amplified from pcDNA3.1-Keima-SKL-Neo and cloned into the *Drosophila* expression vector pUAST.attB to generate the pUAST.attb-Keima-SKL construct [16]. Both pUAST.attB-EGFP-mCherry-SKL (pexo-QC) and pUAST.attb-Keima-SKL (pexo-Keima) constructs were verified by sequencing before sending for phiC31-mediated transgenesis at the *Drosophila* Research facility, University of Manchester. Plasmids were integrated into attP40 sites. Several transformant lines were examined for consistency before settling on a single line for future investigation.

### mRNA extraction from whole Drosophila and RT-qPCR analysis

mRNA, was extracted from whole *Drosophila melanogaster* using the TRIzol method as showed previously (Bogart & Andrews, 2006). Briefly, 50 mg adult fly’s heads or 50 mg L3 Larvae whole body were collected and homogenised in 1 mL of TRIzol reagent (Thermo Fischer Scientific, #15596026-100ML). After adding chloroform, the mixture was centrifuged at 12,000 × g for 15 minutes at 4°C to separate the RNA-containing aqueous phase from the organic phase. The RNA was precipitated from the aqueous phase using isopropanol, washed with 75% ethanol, and resuspended in RNase-free water. The quality and quantity of the extracted mRNA were assessed using a NanoDrop spectrophotometer. The extracted mRNA was kept at -80°C until it was used in subsequent applications like cDNA synthesis and gene expression analysis. RevertAir H Minus reverse transcription (Thermo Scientific, #11541515), RNasin (Promega, #N251S), oligo (dT) 15 primers (Promega, #C1101), and PCR nucleotide mix (Promega, #U144B) were used to synthesise cDNA from 1 mg of RNA. Quantitative PCRs were conducted in triplicate using primers against acbd4-5 (Pair 1; 5’- CGCGGATGCTTCAGTCCAAT-3’; 5’- CTCGTGCCCGTAACCGAAGAT-3’; Pair 2; 5’- TTTCCTGGTGAGGCGGTTCG -3’; 5’- GCTAAAGACCAAGCTCGTGCC-3’). In a Biorad CFX Connect real-time system, primer and cDNA reactions were executed using iTaq Mastermix (Bio-Rad, #172-5171). Ct values (mean cycle threshold) were normalised to tubulin, raised to the exponent of 2^-ΔΔCt^ and normalised to the appropriate control fly line.

### Lysotracker assay

*Drosophila* larval epidermis was dissected in PBS and incubated for 10 minutes with 0.7 mM LysoTracker Deep Red (Invitrogen). Dissected tissues were immersed in PBS on glass slides, covered, and imaged immediately with a 3i spinning disk confocal fluorescence microscope (Intelligent Imaging Innovations, Germany, SlideBook 3i v3.0) equipped with a 63x/NA 1.4 oil objective and a sCMOS camera, Hamamatsu. The protocol was adapted from Lee et al.[18].

### Drosophila pexo-QC embryo preparation and imaging

The protocol was adapted from previous publications [49, 50]. Late stage 16 embryos of the desired genotypes were collected at 22°C and dechorionated by incubating in a solution of 50% bleach and 50% water for 1.5 minutes, followed by extensive rinsing in water to stop the reaction. The dechorionated embryos were then washed in ddH_2_O and spread on a fresh 2% agar/apple juice plate. Late stage pexo-QC-positive (Green+/Red+) embryos were harvested using a Leica MZ10F-FLUO fluorescent microscope. Only embryos lacking the chorion (no dorsal appendices) and with an intact vitellin outer layer were selected. After collection, the pexo-QC embryos were immediately transferred onto an embryo chamber, consisting of a coverslip attached to double-sided sticky tape forming a channel filled with halocarbon oil 700 (Sigma; H8898-50ML) [49]. The living embryos were spaced appropriately within the channel and covered by another coverslip before being imaged using a spinning disk confocal microscope (3i, Intelligent Imaging Innovations, Germany, SlideBook 3i v3.0) equipped with a 10x NA 0.3 objective lens and a sCMOS camera, Hamamatsu.

### Light sheet imaging and sample preparation of whole body Drosophila larvae and white prepupae

Digestion and fixation steps of both 3^rd^ instar larvae and white prepupae were adapted from Pende et al. [51]. Briefly, samples were treated at 37°C for 45 minutes with 0.03% proteinase K (Sigma, #P2308-10MG) in pre-warmed PBS to promote digestion of the superficial layer of larvae and prepupae. After 3x washes in PBS at room temperature for 10 minutes, samples were fixed in 4% paraformaldehyde at 4°C overnight followed by 3x washing with PBS at room temperature for 30 minutes. After the washing steps, animals were immediately mounted on a glass capillary tube (Brand GmbH, Wertheim, Germany) for Light sheet imaging. To do that, 1% Agarose (w/v; Agarose low melt, Roth, Germany) was melted in a heat block at 82°C. The melted agarose was allowed to cool to ∼50°C. A drop was poured on the *Drosophila* samples and the mixture was sucked into a glass capillary tube, allowed to solidify, and visualised in a Z.1 Light-Sheet microscope (Zeiss, Germany) with a 5X/0.1 illumination objective and an EC Plan Neofluar 5X/0.16 detection objective, using Zen software (Zeiss, Germany) for image acquisition and processing. Images were acquired in Multiview with 0° and 180° angles (ventral and dorsal sample side) using a pco.edge scientific complementary metal-oxidase-semiconductor (sCMOS) camera (PCO, Germany). Images acquired using two-side illumination were automatically fused. For each Multiview, a Z-stack maximum intensity projection was generated using Fiji v2.9.0 (ImageJ) [52]. The light sheet analysis was performed for three independent experiments.

### Colocalisation of peroxisomal markers

Dissected pexo-QC larval epidermis sample expressing Acbd4-5-3xHA were fixed using 4% paraformaldehyde in PBS, permeabilised with 0.2% Triton X-100 in PBS, stained with anti-HA primary (Covance, #MMS-101P, 1:50) and AlexaFluor-647-coupled secondary antibodies. For pexo-QC (EGFP channel) and PMP34-Cer co-localisation in the larval epidermis, the tissues were imaged immediately after dissection and samples were imaged using a Zeiss LSM900 with Airyscan (63x/NA 1.4 oil; Zen Blue software). For pexo-Keima (mKeima-SKL) samples, colocalisation were performed with dissected unfixed larval epidermis using a 3i Marianas spinning disk confocal microscope (63x/ NA 1.4, Hamamatsu sCMOS camera; Slide Book 3i v3.0 Software).

### Pexophagy analysis

For *in vivo* nervous system pexophagy studies, L3 larvae were immobilised in halocarbon oil containing 10% chloroform and mounted on a glass slide with a top coverslip (Figure 4A). Adult *Drosophila* were immobilised on ice. Thereafter, brains were carefully dissected in Dulbecco’s PBS (Sigma, RNBF2227) utilising Dumont No. 5 Forceps (Dumont No.5, #11295-10) and delicately placed in MatTek 35mm dishes (MatTek, #P35G-1.5-14-C) filled with PBS. A coverslip was firmly attached to the dish using double-sided tape. *In/ex vivo Drosophila* sample images were acquired using a 3i Marianas spinning disk confocal microscope (10x air NA 0.3, 40x oil NA 1.3 and 63x oil NA 1.4 objectives, Hamamatsu sCMOS camera; Slide Book 3i v3.0 Software, 25°C incubator). Images were acquired sequentially using the following settings: pexo-QC samples: 488 nm laser, 525/30; 561 nm laser, 617/73 nm emission, Keima-SKL samples 445 nm laser, 617/73 nm emission; 561 nm laser, 617/73 nm emission. Determination of pexophagy levels in pexo-QC samples was performed using the semi-automated “mito-QC Counter” plugin implemented in Fiji v2.9.0 software as previously described [53]. The analysis of pexophagy involved three independent experiments, with at least three animals per condition acquired in each experiment.

### Statistical analysis

P-values are calculated using unpaired t-test, one-way ANOVA and Dunnett’s multiple comparisons post hoc test and are denoted as * *P* < 0.05, ** *P* < 0.01, *** *P* < 0.001 and **** *P* < 0.0001. GraphPad Prism 9 was used for all statistical analyses.

## Supplementary Figures

**Figure S1:**
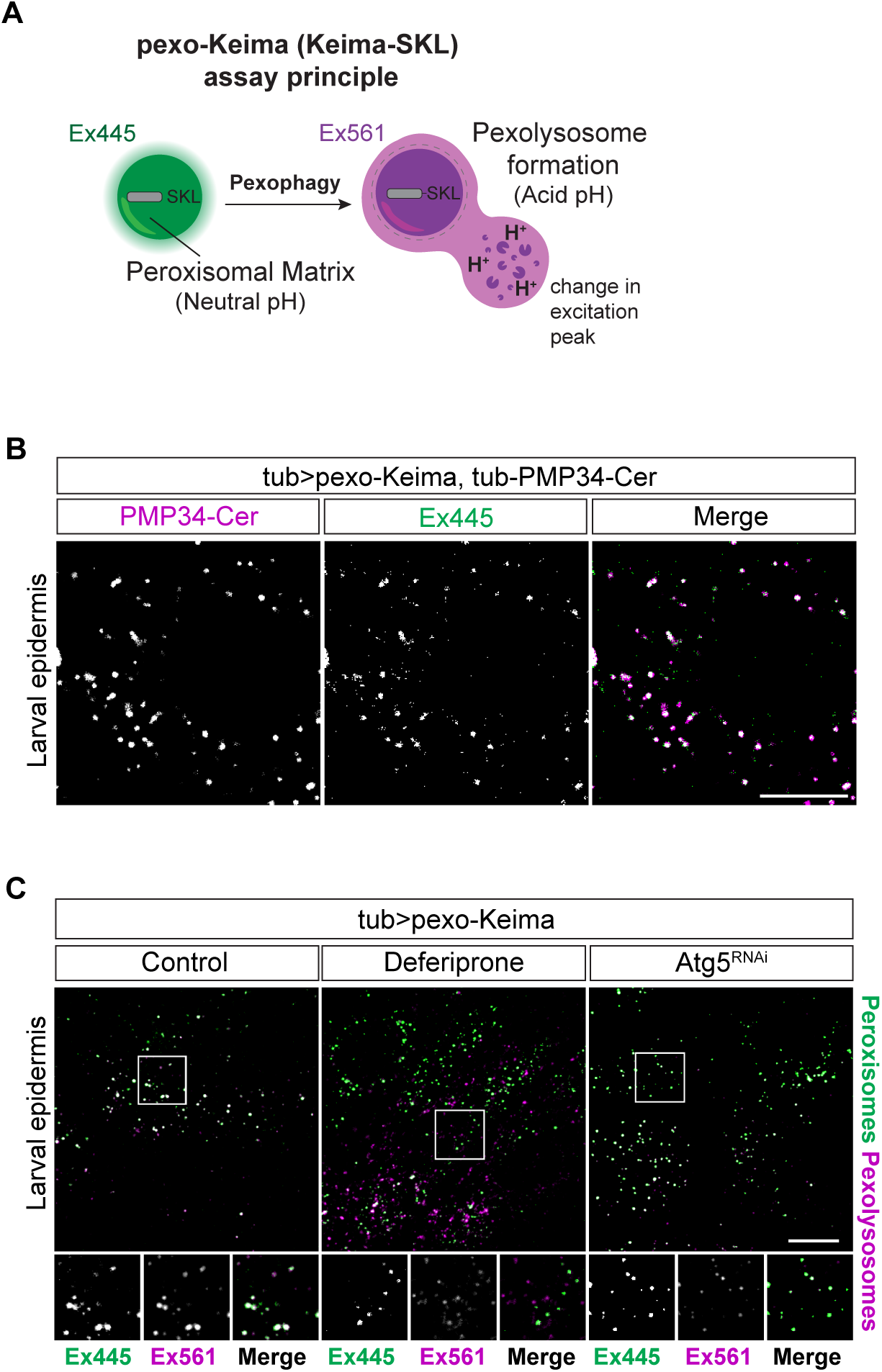
Characterisation of the pexo-Keima (mKeima-SKL) *Drosophila* melanogaster. (**A**) Schematic of the pexo-Keima (mKeima-SKL) pexophagy reporter system. The excitation spectrum of peroxisomal matrix targeted Keima is red shifted upon delivery to the acidic environment of lysosomes, forming pexolysosomes. Images acquired using two excitation wavelengths, 445 nm and 561 nm, and a unique Emission bandwidth (617-673 nm) are represented in green and magenta pseudocolours, respectively. (**B**) Confocal live imaging of 3^rd^ instar larva epidermis comparing the Ex445 fluorescence signal (green) of pexo-Keima with exogenous expression of peroxisomal membrane protein PMP34-Cer (tub-PMP34-Cerulean, magenta). Representative of three independent experiments, with at least three animals analysed per condition in each experiment. (**C**) Comparison of pexolysosome levels in pexo-Keima larval epidermal cells upon exposure to the iron chelator deferiprone (DFP, 65 *µ*M) or Atg5 depletion (Atg5^RNAi^). Representative of two independent experiments, with at least three animals analysed per condition in each experiment. Scale bars: 10 μm. Genotypes analysed were *w^1118^; UAS-Keima-SKL/+; tub-Gal4/tub-PMP34-Cer* (B), *w^1118^; UAS-Keima-SKL/+; tub-Gal4/+* (C), and *w^1118^; UAS-Keima-SKL/+; tub-Gal4/+, w^1118^; UAS-Keima-SKL/UAS-Atg5^RNAi^; tub-Gal4/+* (D).

**Figure S2:**
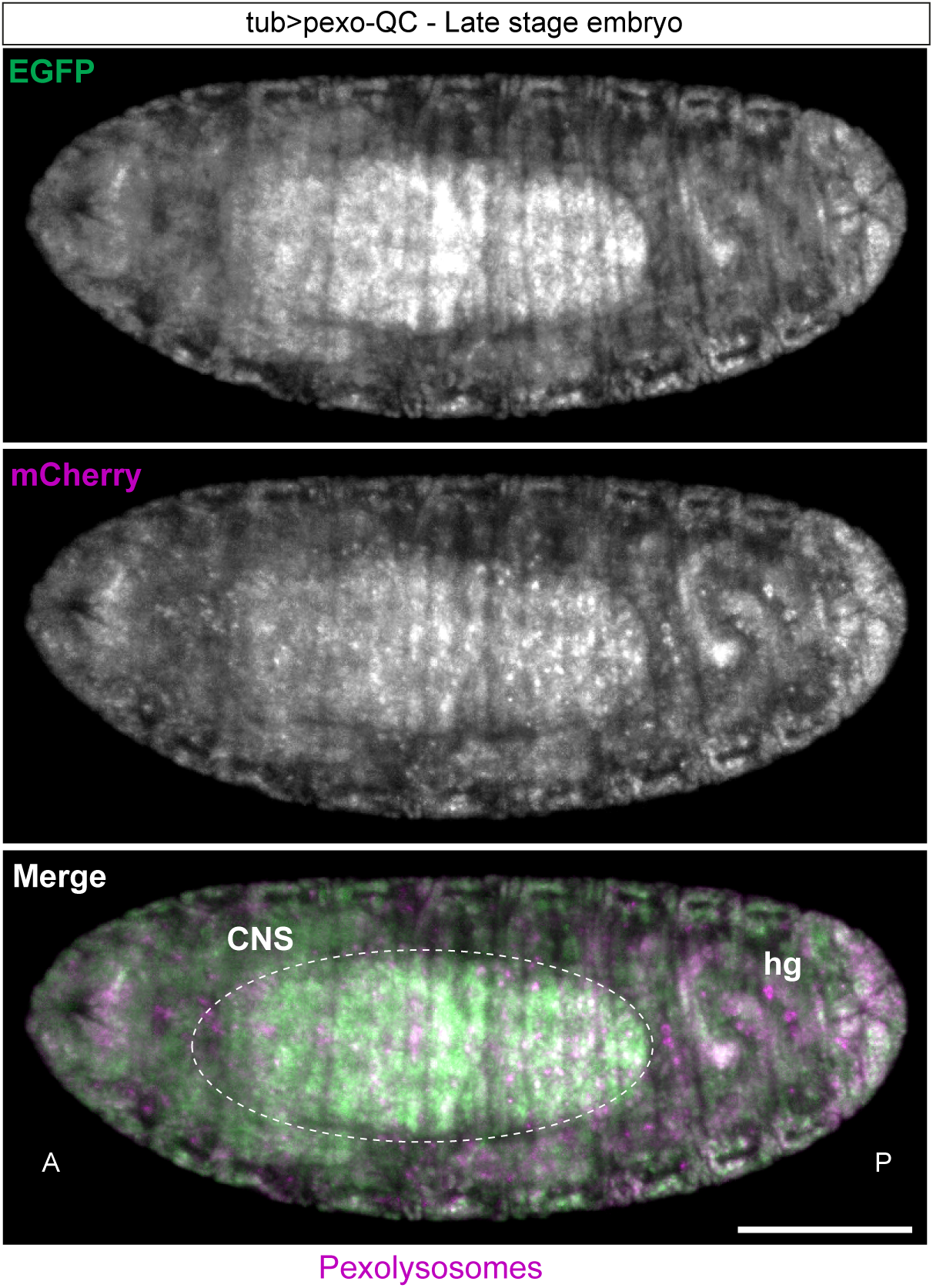
Pexophagy analysis in late stage 16 living pexo-QC embryo. Representative confocal z-stack maximum intensity projection images of the late-stage 16 pexo-QC embryo, acquired from a ventral view with a 10x air NA 0.3 objective. Pexophagy (mCherry, magenta) is distributed spatially across tissues. Central Nervous System (CNS); hg, hind gut. Scale bar: 100 μm. Genotype analysed was w^1118^; UAS-pexo-QC/+; tub-Gal4/+.

**Figure S3:**
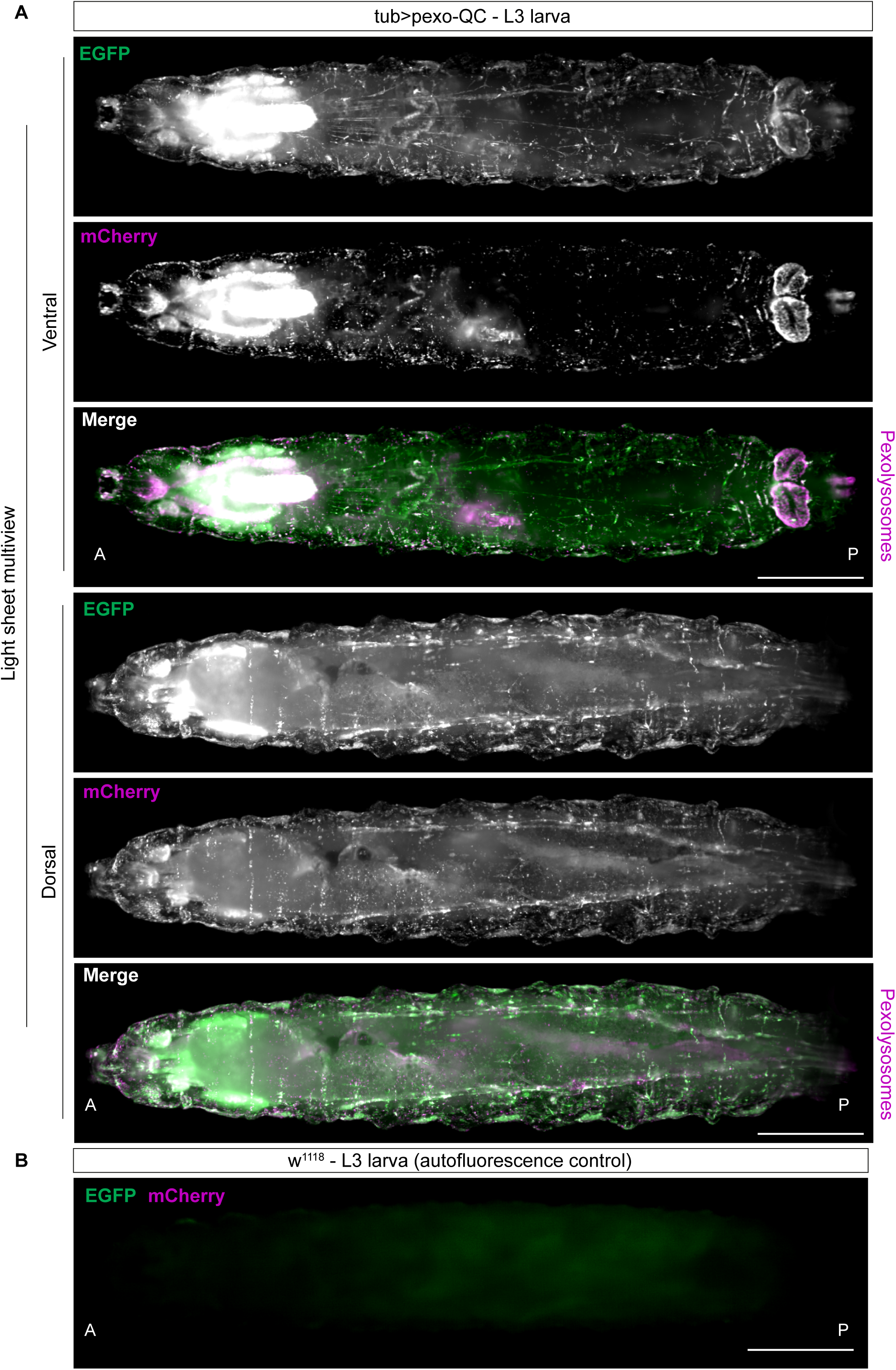
Pexophagy in light sheet pexo-QC larvae. (**A**) Expanded view of Light Sheet Fluorescence Microscopy (LSFM) Multiview images of L3 pexo-QC larva (tub>pexo-QC) shown in Figure 2, providing both single channel images (EGFP and mCherry). (**B**) Panel shows the background autofluorescence observed for the corresponding w^1118^ larva that does not express pexo-QC. P: Posterior; A: Anterior. Scale Bars: 500 μm. Genotype analysed was *w^1118^; UAS-pexo-QC/+; tub-Gal4/+ (A), w^1118^* (B).

**Figure S4:**
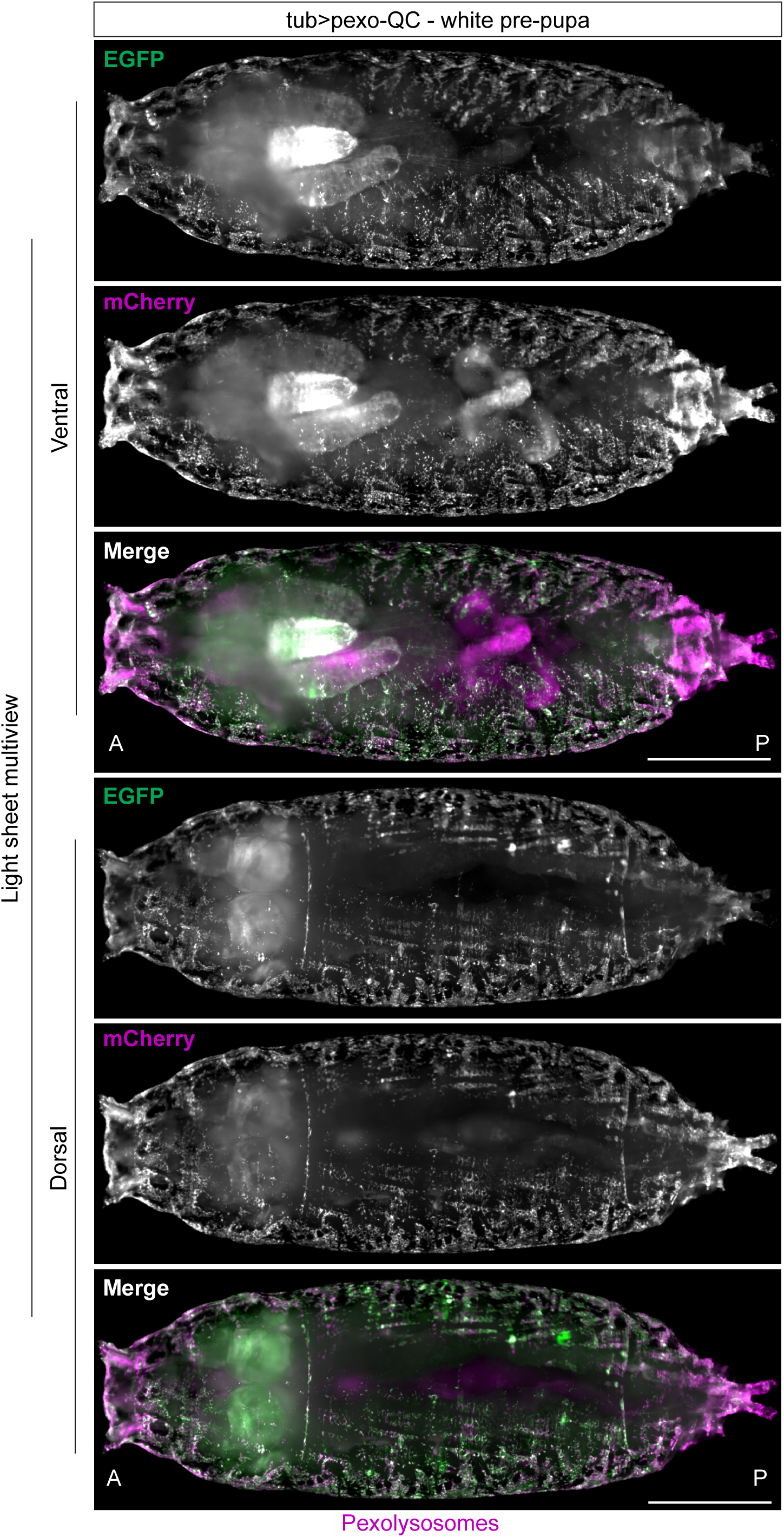
Pexophagy in light sheet pexo-QC white pre-pupa. Expanded view of light sheet fluorescence microscopy (LSFM) Multiview images of pexo-QC white pre-pupa (tub-Gal4>pexo-QC) shown in Figure 2, detailing both single channel images (EGFP and mCherry). Scale Bars: 500 μm. Genotype analysed was *w^1118^; UAS-pexo-QC/+; tub-Gal4/+*.

**Figure S5:**
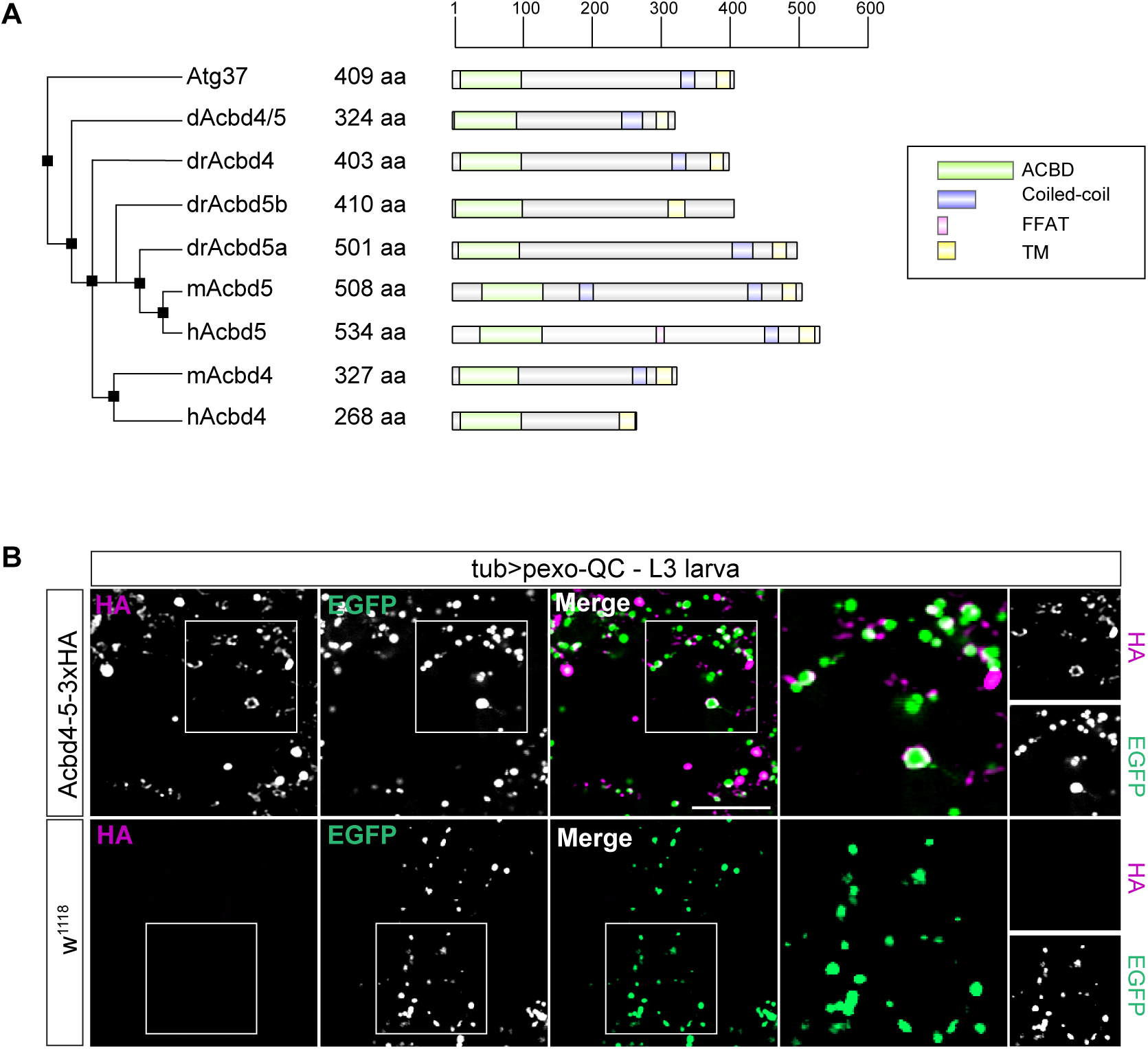
CG8814, the *Drosophila* homolog of human Acbd4-5 localises to peroxisomes. (**A**) The phylogenetic tree and domain structures of *Drosophila* Acbd4-5 (CG8814) and its orthologs in yeast (Atg37), zebrafish (drAcbd4, drAcbd5a, and drAcbd5b), mice (mAcbd4 and mAcbd5), and humans (hACBD4 and hACBD5) are depicted in a schematic image. Domains were annotated in IBS based on InterPro [54]. The phylogenetic tree was generated using Jalview [55]. The ACBD motifs (green), coiled-coil (CC, purple), the transmembrane domain (TM, yellow) and FFAT (pink) motifs are shown. (**B**) Top: Representative high resolution Airyscan images of dissected and fixed epidermis of L3 larvae expressing pexo-QC and Acbd4-5-3xHA (tub>pexo-QC;tub>Acbd4-5-3xHA) stained with HA antibody. Bottom: Negative control sample, using the same experimental setup, but pexo-QC larvae were crossed with w^1118^ flies, lacking expression of Acbd4-5-3xHA (negative control). Scale bar: 10 μm. Genotypes analysed were *w^1118^; UAS-pexo-QC/+; tub-Gal4/+, w^1118^; UAS-pexo-QC/UAS-Acbd4-5-3xHA; tub-Gal4/+*.

**Figure S6:**
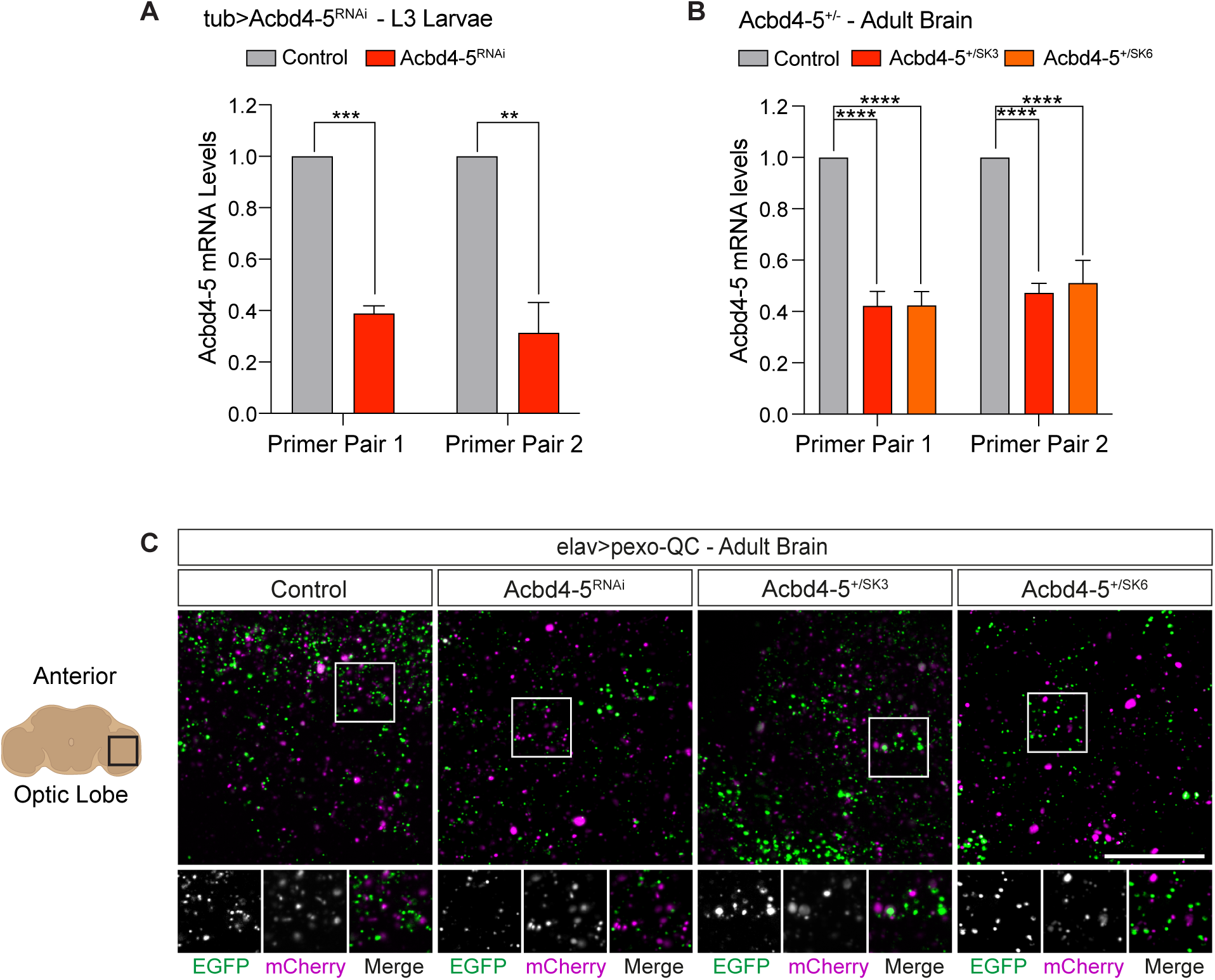
Acbd4-5 mRNA levels and pexophagy in the larvae and adult brain of Acbd4-5 ablated *Drosophila*. (**A-B**) Quantitative RT-PCR reactions for Acbd4-5 mRNA (normalised to tubulin) were performed with cDNA derived from control and tub>Acbd4-5^RNAi^ L3 Larvae and tub>Acbd4-5RNAi, Acbd4-5^+/SK3^ and Acbd4-5^+/SK6^ KO fly heads. Two primer pairs were used per target from three independent experiments. Error bars show SD of the fold change (2^-ΔΔCt^) of Acbd4-5 mRNA in each heterozygote Acbd4-5 combination, normalised to their matched control flies. Statistical significance determined by unpaired t-test (A) and one-way ANOVA with Dunnett’s multiple comparison test (**B**); \*\**P* < 0.01, \*\*\**P* < 0.001 and \*\*\*\**P* < 0.0001. Supplementary to Figure 6 A-C. (**C**) Representative Z-stack maximum intensity projection confocal images of *ex vivo* adult brain (9-12 days old) neurons, expressing pexo-QC ± Acbd4-5^RNAi^ or Acbd4-5^+/SK3^ or Acbd4-5^+/SK6^. Scale bars: 20 μm. Supplementary to Figure 6 G-H. Genotypes analysed were *w^1118^; UAS-pexo-QC/+; tub-Gal4/+, w^1118^; UAS-pexo-QC/UAS-Acbd4-5^RNAi^; tub-Gal4/+* (A), *w^1118^; UAS-pexo-QC/+; elav-Gal4/+, w^1118^; UAS-pexo-QC/ Acbd4-5^SK3^; elav-Gal4/+, w^1118^; UAS-pexo-QC/Acbd4-5^SK6^; elav-Gal4/+* (B), *w^1118^; UAS-pexo-QC/ +; elav-Gal4/+, w^1118^; UAS-pexo-QC/UAS-Acbd4-5^RNAi^; elav-Gal4/+; UAS-pexo-QC/Acbd4-5^SK3^; elav-Gal4/+, w^1118^; UAS-pexo-QC/Acbd4-5^SK6^; elav-Gal4/+ (C)*.

